# ISO-Relevance Functions - A Systematic Approach to Ranking Genomic Features by Differential Effect Size

**DOI:** 10.1101/381814

**Authors:** Soumyashant Nayak, Nicholas F. Lahens, Eun Ji Kim, Emanuela Ricciotti, George Paschos, Sarah Tishkoff, Dimitra Sarantopoulou, Shaon Sengupta, Barry Cooperman, Tilo Grosser, Gregory R. Grant

**Affiliations:** Perelman School of Medicine, University of Pennsylvania, Philadelphia, PA 19104

## Abstract

It is common to measure a large number of features in parallel to identify those differing between two experimental conditions - e.g. the search for differentially expressed genes using microarrays or RNA-Seq. Ranking features by *p*-value allows for control of the TYPE I error, but *p*-values are not reliable when there are very few replicates; and investigators typically require features be ranked by “fold change” 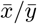 in conjunction with *p*-values. At first glance the fold change appears to be a natural quantity on which to compare the differential behavior of features. But it is highly sensitive to small values in the denominator and is problematic in how it equates changes in both small and large numbers such as a change from 1 to 2 versus a change from 100 to 200. The strategy of adjusting all values by adding one is a widely used heuristic approach to try to mitigate the problems with fold-change. However, that can be far from optimal. A systematic strategy to determine an optimal value (pseudocount) to adjust by is employed using both real and simulated benchmark data. In RNA-Seq a value of 20 appears to be close to optimal in all cases. Another strategy is to sort by difference 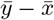, but this is problematic for comparing measurements across a wide spectum, as large differences of small values rank below proportionally smaller difference in large values. An abstract mathematical framework is introduced to describe the problem of ranking by differential effect size, enabling us to study the ranking problem in general as opposed to specific contexts such as fold-change or difference. From this framework we discovered a remarkable property of pseudocounts, in that they strike a balance between sorting by fold-change and sorting by difference. Lastly, another fundamentally different type of application is presented, which is to rank di-codons by their differential abundance in the ORFeome of different species.

## 1. Introduction

A canonical example to illustrate the problem at hand is that of differential gene expression analysis using (normalized) RNA-Seq read counts. The goal is to find the genes whose expression is “different” between two populations of samples. To do this researchers generate measurements for tens of thousands of genes in a number of replicate samples from each of the two conditions. Typically *p*-values or *q*-values are calculated for each gene and then genes are ranked by these values. In this way the genes which are most differential are at the top, sorted by the power of the statistical evidence for differential expression. A secondary measure, and often the primary measure, used to rank genes by their differential effect is fold-change. One might argue that one should not be ranking features by effect sizes like fold-change and should instead always use statistically rigorous metrics such as *p*-values. However, investigators often want to rank features by fold change as a secondary ranking in conjunction with *p*-values, because of granularity of the *p*-values. For example if the top significance level by *p*-value has 1000 genes all equally significant, which commonly happens if there are few replicates, it may then be desirable to rank these 1000 genes also by fold change. Additionally, often genes with very low *q*-values also have very low fold-changes like 1.01 fold and such low fold-changes are generally not considered biologically relevant.

It is universally understood that fold-change is problematic when the denominator is zero, so to get around this one typically adds some fixed positive constant, known as a “pseudocount,” to both numerator and denominator. Most of the time this pseudocount is taken to be one. The use of pseudocounts in ranking differential effects is perhaps one of the most widely used heuristics in biology and does mitigate the “division by zero” problem. However the division by zero problem is not the only problem nor the biggest problem with this approach. If the measurements span a wide spectrum of values, fold-change becomes problematic. To illustrate this consider the following example. Suppose the data in question is RNA-Seq data and the measurements are gene level counts of reads. In this type of data counts typically range from zero to hundreds of thousands and even higher. Suppose we have one sample called Treatment and one sample called Control. Consider two hypothetical genes, *G*_1_ and *G*_2_ with the following expression counts:

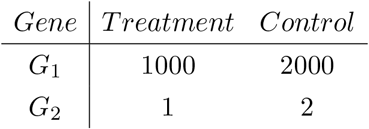

The fold change equals two for both gene *G*_1_ and *G*_2_. However, we would not want to say these genes have equal differential effects. The discrete nature of RNA-Seq only exacerbates this problem. The only possible values are integer counts 0, 1, 2, etc. Therefore a change from 1 to 2 is very likely to be due to random variation. On the other hand a change from 1000 to 2000 is likely to reflect a real difference in population means.

Adding a pseudocount to all values, typically done to solve the division by zero problem, also mitigates this problem. But continuing with the example, if we use a pseudocount of one we would then have

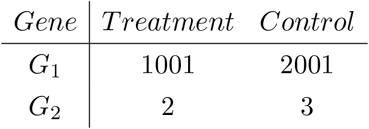

The fold-change for *G*_2_ went from 2 without the pseudocount, to 1.5 with, while the fold-change for *G*_1_ went from 2 to 1.999. However *G*_2_ would still have the same fold change as *G*_1_ which went from 1000 to 1500 and this is still far from desirable. Yet virtually all studies that employ a pseudocount use a pseudocount of one. There is no reason, however, to take a purely heuristic approach to this problem. Using carefully designed benchmarking data as well as real data, it is possible to perform a systematic analysis to optimize pseudocounts. We do this below in Results.

Another natural method to rank the differential behavior of features is to use difference instead of fold change. This eliminates the issue of dividing by zero; however a different problem arises, which is illustrated by the following example:

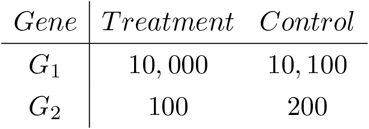

In this case both genes have a difference of 100 while we would definitely want to prioritize *G*_2_ over *G*_1_. We see that just as fold-change has problems with small values (and particularly with zero), difference has problems with large values. We prove below that pseudocounts provide a flexible balance between the two extremes of fold-change and difference.

These considerations lead naturally to a general theoretical framework that systematically describes the concept of ranking features. In order to say two features have different differential behavior from each other, we first need to develop the concept of changes being equally meaningful. This leads to the definition of “iso-relevance functions”. The general framework is described first, followed by applications to RNA-Seq differential gene expression analysis and differential di-codon usage between species.

It is helpful to keep in mind the differential gene expression example, however we will discuss another application of these methods to rank di-codons by their differential use in two species, where there are no distributions or a concept of the underlying truth. The goal in that case is simply to prioritize features by their ability to perform a downstream task. The concepts described below are relevant to a wide range of applications.

There will always be some amount of subjectivity in deciding whether two differential effects are equally significant or not. However that’s not to say it is arbitrary. Consider a gene which changes from 1000 to 2000 in RNA-Seq counts. Anybody would deem a change from 0 to 1 to be less significant than that. Similarly a change from 0 to 100 would probably be deemed more significant than 1000 to 2000. Somewhere in between is a grey area and we will see that the concept of pseudocount is tied implicitly, but intimately, with making a judgement call in this grey area. The grey area will rely on the intuition of the investigator, however the ability to systematically quantify that intuition is what is under discussion here.

## 2. Mathematical Framework

In this section, we first illustrate how ranking by fold-change and ranking by difference are manifestations of the same ranking measure but at different scales. We state this as Main Result 1 below. After proving it, we develop the concept of iso-relevance functions as the appropriate framework for a general theory of ranking measures which yields an interpretation of the pseudocount based on a linear (complete) family of iso-relevance functions.

### Main Result 1

*Consider n-tuples*, (*x*_1_,*x*_2_,…, *x_n_*) *and* (*y*_1_, *y*_2_,…, *y_n_*) *representing measurements for features G*_1_, *G*_2_, …, *G_n_ under conditions x and y respectively. Then there is a C* > 0 *such that for all a* > *C the ordering of the features obtained using differences x_i_* – *y_i_, is the same as the ordering obtained using adjusted fold-changes* 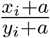.

This follows from the following proposition and its corollary.

### Proposition 2.1.

*Consider four positive real numbers x*_1_, *y*_1_, *x*_2_, *y*_2_ *such that y*_1_ – *x*_1_ > *y*_2_ – *x*_2_. *There is a non-negative real number A such that for all a* > *A*,

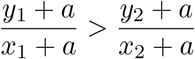

*Proof*. Note that (*y*_1_ – *x*_1_) – (*y*_2_ – *x*_2_) is a strictly positive real number. Also we have the following algebraic identity.

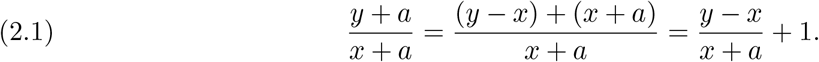

Let *a* be a positive real number. From (2.1) it follows that

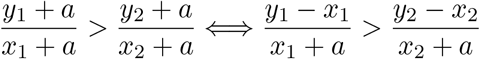

Keeping in mind that *x*_1_ + *a*, *x*_2_ + *a* are positive numbers, after cross-multiplication, we have that,

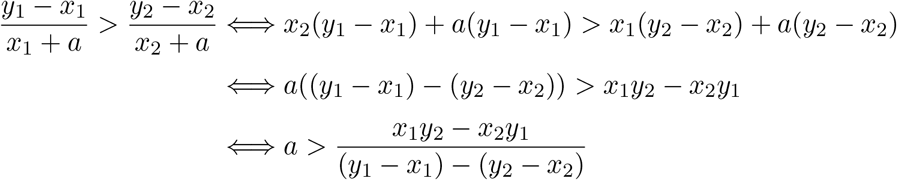

Thus we have shown

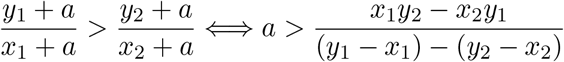

Thus the proposition is proved by setting

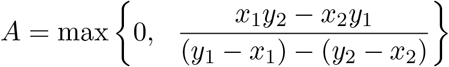

### Corollary 2.2.

*Let* (*x*_1_, *x*_2_,…, *x_n_*) *and* (*y*_1_, *y*_2_,…, *y_n_*) *be n-tuples of positive real numbers such that y*_1_ – *x*_1_ > *y*_2_ – *x*_2_ > ··· > *y_n_* – *x_n_*. *There is a non-negative real number A such that for all a* > *A*,

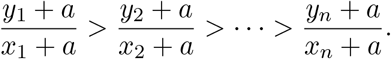

*Proof*. Let

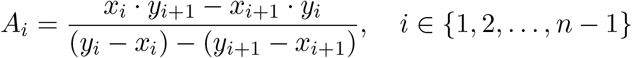

By the previous proposition, for *a* > *A_i_*, we have that,

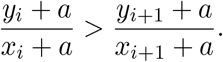

Thus if we define

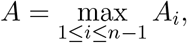

then for all *a* > *A* it follows that

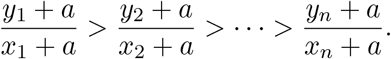

Let ℝ_+_ denote the set {*x* ∈ ℝ | *x* ≥ 0}. Suppose we have *n* features *f*_1_,…,*f_n_* and measurements *x*(*f_i_*) ∈ ℝ_+_ and *y*(*f_i_*) ∈ ℝ_+_ for each *i* = 1,…, *n*. In light of the above results we see that the sorting features by the ratio

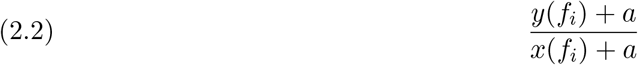

is the same as sorting by fold-change *y*(*f_i_*)/*x*(*f_i_*) when *a* = 0 and is the same as sorting by difference *y*(*f_i_*) – *x*(*f_i_*) when *a* is sufficiently large. Let *C* be the infimum of all values with the property that sorting using a pseudocount of *a* > *C* is equivalent to sorting by difference. Suppose 0 < *a* < *C*, then sorting by (2.2) with this value of *a* strikes a balance between fold-change and difference. The closer *a* is to zero the more weight fold-change has in this balance, and the closer *a* is to *C* the more weight difference has in this balance. Once *a* > *C* then difference becomes 100% of the balance.

Each particular application, e.g. a specific RNA-Seq differential expression analysis, has optimal values of *a* that strike the balance between fold change and difference that achieves the most accurate ranking possible using this approach. Such optimization can be done with benchmark data, when available. If we can find a range of values of *a* that always tend to be more optimal than another range, this information can then be useful in practice. For example, suppose we find that any value of *a* in the interval [10, 30] is almost always better than any value of *a* in the interval [0, 5] then we can use this information to give some guidance. There’s no reason to expect this to be possible *a priori* however as will be shown below, in RNA-Seq analysis the optimal value of *a* appears to be remarkably stable at around *a* = 20.

In the rest of this section, we develop the appropriate mathematical framework to study ranking measures.

### Definition 2.3.

We say that a function *f* : ℝ_+_ → ℝ_+_ is an *iso-relevance function* if the change *x* → *f*(*x*) is (deemed) equally relevant for all *x* in ℝ_+_.

A straightforward example of an iso-relevance function is *f*^(0)^(*x*) = *x* which stands for zero change.

### Definition 2.4.

Let Ω = {(*x, y*) ∈ ℝ^2^ | *x* ∈ ℝ_+_, *y* ≥ *x*} ⊂ ℝ_+_ × ℝ_+_, we refer to Ω as the *relevance octant*.

### 2.1. What properties should an iso-relevance function *f* (≠ *f*^(0)^) have?

One iso-relevance function should correspond to one effect size, e.g. two-fold change. We note some properties one may reasonably expect of such a function; followed by justifications.

P0. *f*(0) ≠ 0,
P1. *f* > *f*^(0)^,
P2. *f* – *f*^(0)^ is strictly increasing,
P3. *f*/*f*^(0)^ is strictly decreasing; 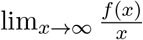 exists and is strictly greater than 1.
P4. *f* is continuous.

Rationales for the above properties follow.

P0. If we think of 0 → 0 as *no change*, then the only iso-relevance function *f* such that *f*(0) = 0 should be given by *f*^(0)^ which corresponds to the null change *x* → *x*.
P1. In this framework, we compare relevance of changes in the same direction (comparing magnitude of changes in different directions can be done by just ignoring the directionality). As (*f* – *f*^(0)^)(0) = *f*(0) ∈ ℝ_+_ is strictly positive, *f*(*x*) – *f*^(0)^(*x*) must be strictly positive for all *x*, otherwise the directionality of the change is different between 0 and some value of *x* ≠ 0, contradicting our assertion of equal relevance for the change at all points in ℝ_+_. Also the change *x* → *f*(*x*) should be equally relevant to the change 0 → *f*(0), and there cannot be multiple choices for *f*(*x*).
P2. For example consider the change 0 → 20 as more relevant than the change 100 → 120 which in turn would be considered more relevant than the change 200 → 220, etc. The same absolute difference (20 in this case) becomes less convincing at higher values. Thus for an iso-relevance function *f*, *f* – *f*^(0)^ should be strictly increasing.
P3. In a similar vein as (ii), we would consider the change 100 → 125 as less relevant than the change 500 → 625 which in turn would be considered less relevant than the change 1000 → 1250. The same relative change (25 % in this case) becomes more convincing at higher values. Thus for an iso-signficance function *f*, *f*/*f*^(0)^ should be strictly decreasing and since it is bounded below by 1, 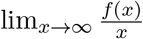 exists. Further we do not want the behaviour of *f*(*x*) to be similar to *x* near ‘infinity’ (at large values) which is why we want to require that 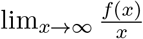 be strictly greater than 1.
P4. Small changes in *x* should lead to small changes in *f*(*x*).

The above defines an iso-relevance function corresponding to a particular level of differential effect size. We now define the notion of a *complete* family of iso-relevance functions corresponding to a continuum of differential effect sizes. It is with this family that we can rank all features in a differential massively parallel experiment with respect to each other, so that the higher a feature is on the list, the more differential the effect is for that feature. A canonical example to keep in mind is that of differential gene expression between two conditions, as measured with RNA-Seq counts or microarray intensities. In this case “relevant” means “differentially expressed.”

We state the properties first and give rationales below. Essentially a family of iso-relevance functions is a partition of the relevance octant into disjoint curves with certain properties (see Fig. 1 for an example).

**Figure 1.**
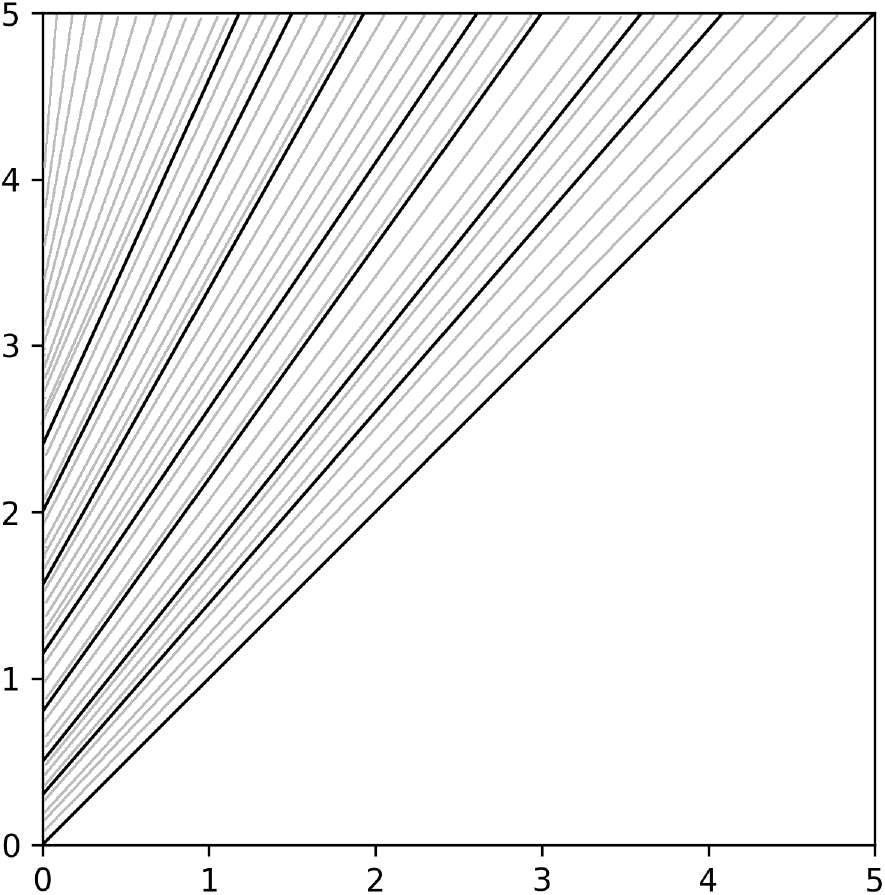
A linear family of iso-relevance functions

### 2.2. What properties should a *complete* family 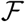 of iso-relevance functions have?

P5. For each *a* > 0, there is a unique 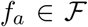 such that *f_a_*(0) = *a*. Consequently, if *f*, 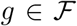 such that *f*(0) = *g*(0), then *f* = *g*,
P6. For any point (*x, y*) in the relevance octant, there is a unique iso-relevance function 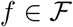 such that *y* = *f*(*x*).
P7. If *f*(0) > *g*(0), then *f*(*x*) > *g*(*x*) for all *x* in ℝ_+_,
P8. If *F* : Ω → ℝ_+_ is defined by *F*(*a, x*) = *f_a_*(*x*) (where *f_a_* is defined as in P5), then the function *F* is continuous,
P9. If *f*, 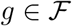, then 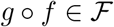.

Rationale for the above properties follow.

P5. Since *a* > 0, for *x* ∈ ℝ_+_, the change 0 → *a* is more relevant than the change *x* → *x* but less relevant than the change *x* → *M*, where *M* denotes an extremely large number (so-called infinity). Thus we expect that there is some value *x*’ > *x* such that 0 → *a* is equally relevant to *x* → *x*’. Defining *f_a_*(*x*) = *x*’, we obtain the desired iso-relevance function. The change *x* → *x*’ and the change *x* → *x*” are equally relevant if and only if *x*’ = *x*”.
P6. For *y* > *x*, the change 0 → 0 is less relevant than the change *x* → *y*, and the change 0 → *y* is more relevant than the change *x* → *y*. Thus there is a number *p* between 0 and *y* such that the change 0 → *p* is equally relevant to the change *x* → *y*. Considering the iso-relevance function *f* which satisfies *f*(0) = *p*, we must have *y* = *f*(*x*). The uniqueness follows from property P5. Thus the iso-relevance functions form a partition of the relevance octant (into disjoint curves).
P7. If *f*(0) > *g*(0), the change 0 → *f*(0) is more relevant than the change 0 → *g*(0). Thus the change *x* → *f*(*x*) (~ 0 → *f*(0)) is more relevant than *x* → *g*(*x*) (~ 0 → *g*(0)).
P8. For *x* in ℝ_+_, the change 0 → *f* (0) ~ *x* → *f*(*x*). Similarly 0 → *g*(0) ~ *x* → *g*(*x*). Thus *x* → *f*(*x*) ~ *x* → *g*(*x*), as *f*(0) = *g*(0). Further, if 0 → *f* (0) is more relevant than 0 → *g*(0), then *x* → *f*(*x*) is more relevant than *x* → *g*(*x*) for all *x* in ℝ_+_.
P9. Let 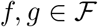. For *x*, *y* in ℝ_+_, the change *x* → *f*(*x*) is equally relevant to the change *y* → *f*(*y*). Similarly the change *f*(*x*) → *g*(*f*(*x*)) is equally relevant to the change *f*(*y*) → *g*(*f*(*y*)). Thus it is not unreasonable to conclude that the change *x* → *g*(*f*(*x*)) (viewed as *x* → *f*(*x*) → *g*(*f*(*x*))) is equally relevant to the change *y* → *g*(*f*(*y*)) (viewed as *y* → *f*(*y*) → *g*(*f*(*y*)). Thus *g* o *f* is also an iso-relevance function.

*Remark* 2.5. If 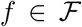, then by repeated application of property P9 one may note that *f*^(*n*)^ = *f* o ··· o *f* (*f* composed with itself n times, *n* ∈ ℕ) is also in 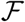.

#### Proposition 2.6.

*A complete family of iso-relevance functions forms a semigroup parametrized by* ℝ_+_, *with the identity element given by f*^(0)^.

*Proof*. From the previous remark, if *f* is an iso-relevance function in 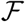, then so is *f*^(*n*)^ for *n* ∈ ℕ. Note that 0 < *f*(0) < *f*(*f*(0)) as *f* is strictly increasing (and *f*(0) > 0 from property P1). Consider the restriction of *F*, defined in property P9, to the diagonal of ℝ_+_ × ℝ_+_. The real-valued function h on ℝ_+_, defined by *h*(*x*) = *F*(*x, x*) is a strictly increasing continuous function as *F* is continuous (property P7). As *h*(0) = *F*(0,0) = 0 and *h*(*f*(0)) = *F*(*f*(0), *f*(0)) = *f*(*f*(0)) > *f*(0), by the intermediate value theorem, we have that *h*(*a*) = *F*(*a, a*) = *f*(0) for some a in (0, *f*(0)). Let *g* be the iso-relevance function such that *g*(0) = *a*. Thus *F*(*g*(0),*g*(0)) = *g*(*g*(0)) = *f*(0). As seen before, *g* o *g* is also an iso-relevance function and (*g* o *g*)(0) = *f*(0), which implies that *g* o *g* = *f*. We denote *g* by *f*^(1/2)^. Repeating the process, we obtain iso-relevance functions of the form *f*^(1/2*n*)^. By composition of these functions, we obtain the iso-relevance functions which can be expressed as (compositional) dyadic powers of *f*.

Note that 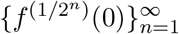 is a strictly decreasing sequence bounded below by 0. Thus it has a limit which we denote by *γ*. Let *ϕ* be an iso-relevance function such that *ϕ*(0) = *γ*. For *n* in ℕ, we have that *ϕ* < *f*^(1/2*n*)^ (property P6). Note that

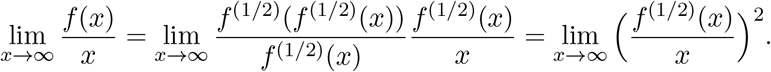

Thus

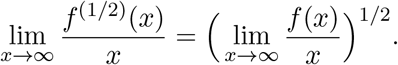

Inductively we can prove that

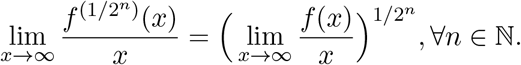

Hence,

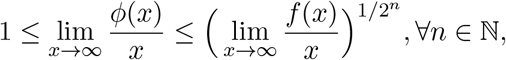

and we conclude that

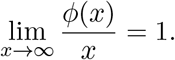

From property P3, we have that *ϕ* = *f*^(0)^ and thus lim_*n*→∞_ *f*^(1/2*n*)^(0) = *γ* = 0.

As dyadic numbers are dense in the reals, using the above conclusion and a routine continuity argument, we can define *f*^(*α*)^ for all positive real numbers *α* such that for *α*, *β* ∈ ℝ_+_, *f*^(*α*)^ o *f*^(*β*)^ = *f*^(*α*+*β*)^. Moreover for any *α* ∈ ℝ_+_ there is an *α* in ℝ_+_ such that *f*^(*α*)^(0) = *a* and *f*^(*α*)^ is the unique iso-relevance function attaining the value *a* at 0.

#### Corollary 2.7.

*A non-trivial iso-relevance function f* (≠ *f*^(0)^) *belongs to precisely one complete family of iso-relevance functions*.

#### Definition 2.8.

In Proposition 2.6, we noted that one non-trivial iso-relevance function in 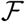 is enough to obtain the rest of the functions in 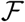. When generating a complete family of iso-relevance functions from one function, we call the generator used the **seed**.

#### Example 2.9.

Let *a, b* be two real numbers such that *a* > 0, *b* > 1. We will see below that the iso-relevance function *f*^(1)^(*x*) = *a* + *bx* is a seed for a complete family of iso-relevance functions. One may interprete the seed in the following manner : the change 0 to *a* is deemed equally relevant to the relative change *x* → *bx* in the limit as *x* → ∞. The other iso-relevance functions in the corresponding complete family are given by

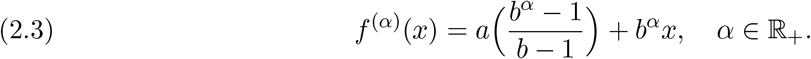

In other words, the change 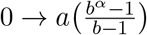 should be deemed equally relevant to the change *x* → *b^α^x* for larger numbers *x*.

In order to see that (2.3) defines a complete family of iso-relevance functions, we only need check properties P6, P7 and P9 as the rest of the properties are straightforward to verify.

<u>Proof of P6.</u> Let (*x,y*) be a point in the relevance octant. Note that

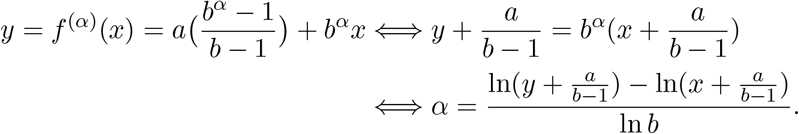

As *y* ≥ *x*, we have *α* ≥ 0 and thus there is a unique iso-relevance function *f*^(*α*)^ such that *y* = *f*^(*α*)^(*x*). <u>Proof of P7</u>. Note that *f*^(*α*)^(0) > *f*^(*β*)^(0) ⇒ *b^α^* > *b^β^* → *α* > *β* ⇒ *f*^(*α*)^(*x*) > *f*^(*β*)^(*x*) for all *x* ∈ ℝ_+_. <u>Proof of P9</u>. For *α*, *β* ∈ ℝ_+_, and *x* in ℝ_+_, we have

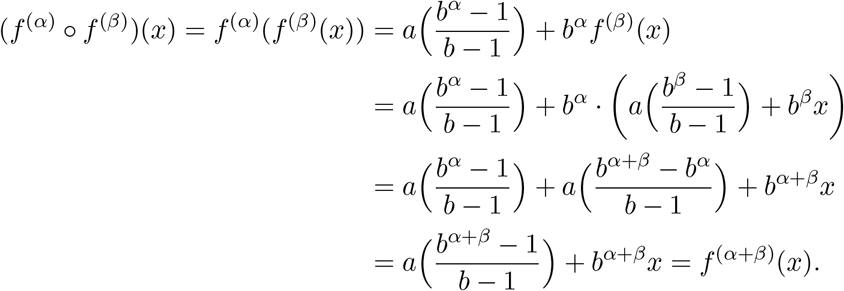

Thus *f*^(*α*)^ o *f*^(*β*)^ is in the family of functions defined.

In Fig. 1, the iso-relevance functions depicted are with respect to the seed given by 2 + 2*x*.

#### Definition 2.10.

Let 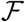 be a complete family of iso-relevance functions with a given seed *f* = *f*^(1)^. For (*x, y*) in the relevance octant, let *α* ∈ ℝ_+_ be such that *y* = *f*^(*α*)^(*x*). Then *α* is called the relevance level of the change *x* → *y* (with respect to 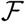).

We state the result in the proof of property P6 in Example 2.9 as a proposition below.

#### Proposition 2.11.

*Let* (*x, y*) *be a point in the relevance octant and consider the complete iso-relevance family generated by a* + *bx where a* > 0, *b* > 1. *Then the relevance level of the change x* → *y is given by*

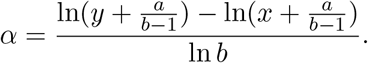

*Remark* 2.12. The above proposition may be interpreted in the following manner. The change *x*_1_ → *y*_1_ is more significant than the change *x*_2_ → *y*_2_ if

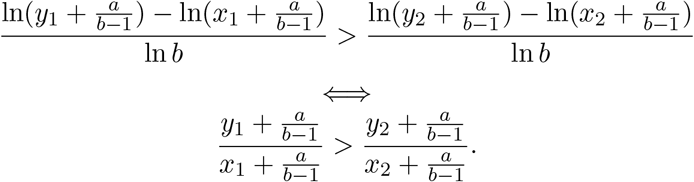

#### Main Result 2

*If we choose b* = 2 *and let a be such that the change* 0 → *a is as significant as the change x* → 2*x (two-fold change) in the limit as x* → ∞, *then* 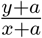 *is a measure of the relevance level of the change. This observation provides a philosophical interpretation of the pseudocount value as the change from* 0 *that is deemed as relevant as two-fold change in the limit*.

## 3. Results

### 3.1. RNA-Seq. Dataset

An investigation into optimal pseudocounts in RNA-Seq differential expresssion (DE) analysis was performed based on twenty RNA-Seq benchmark datasets, obtained from transformations of real data, from several different organisms and tissues:

- Human Blood (≈ 9 million reads)
- Human UHRR (≈ 9.5 million reads)
- Mouse Liver (≈ 21 million reads)
- Mouse Aorta (≈ 37 million reads)
- Arabidopsis (≈ 23 million reads)

Details on how the benchmarking data was generated are given in Materials and Methods. Essentially these are real data with all the properties of real data, with some number of differentially expressed genes created by random subsampling of reads. Each dataset has two conditions of 8 replicates each. In half of the datasets, there are 200 differentially expressed genes and in the other half there are 1000. These are divided into two types; Balanced datasets (B) have half DE genes upregulated and the other half downregulated, and unbalanced datasets (U) have all genes upregulated. For each dataset a graph was prepared that plots the pseudocount c on the horizontal axis against the size of the intersection of the top *N* genes in the list sorted by 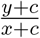 to the known DE genes list where *y* is the average of the gene’s expression values over the eight replicates in the first group, and *x* is the average of the gene expression values over the eight replicates in the second group, and *N* is either 200 or 1000 depending on whether the data set has 200 or 1000 DE genes. The results for the 200 DE genes datasets (1000 DE genes datasets, respectively) are shown in Fig. 2 (Fig. 3, respectively).

**Figure 2.**
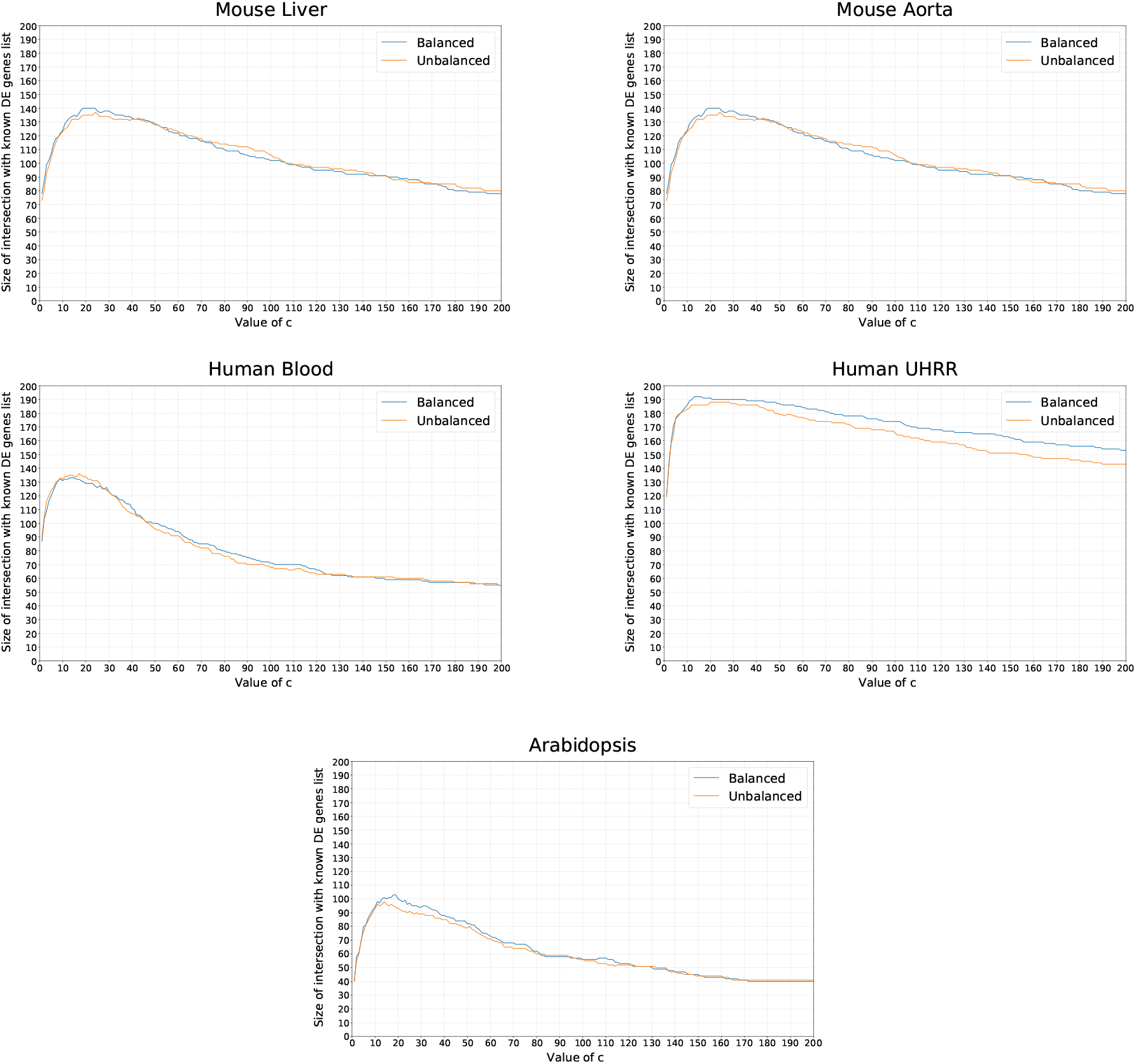
Results for datasets with 200 DE genes

**Figure 3.**
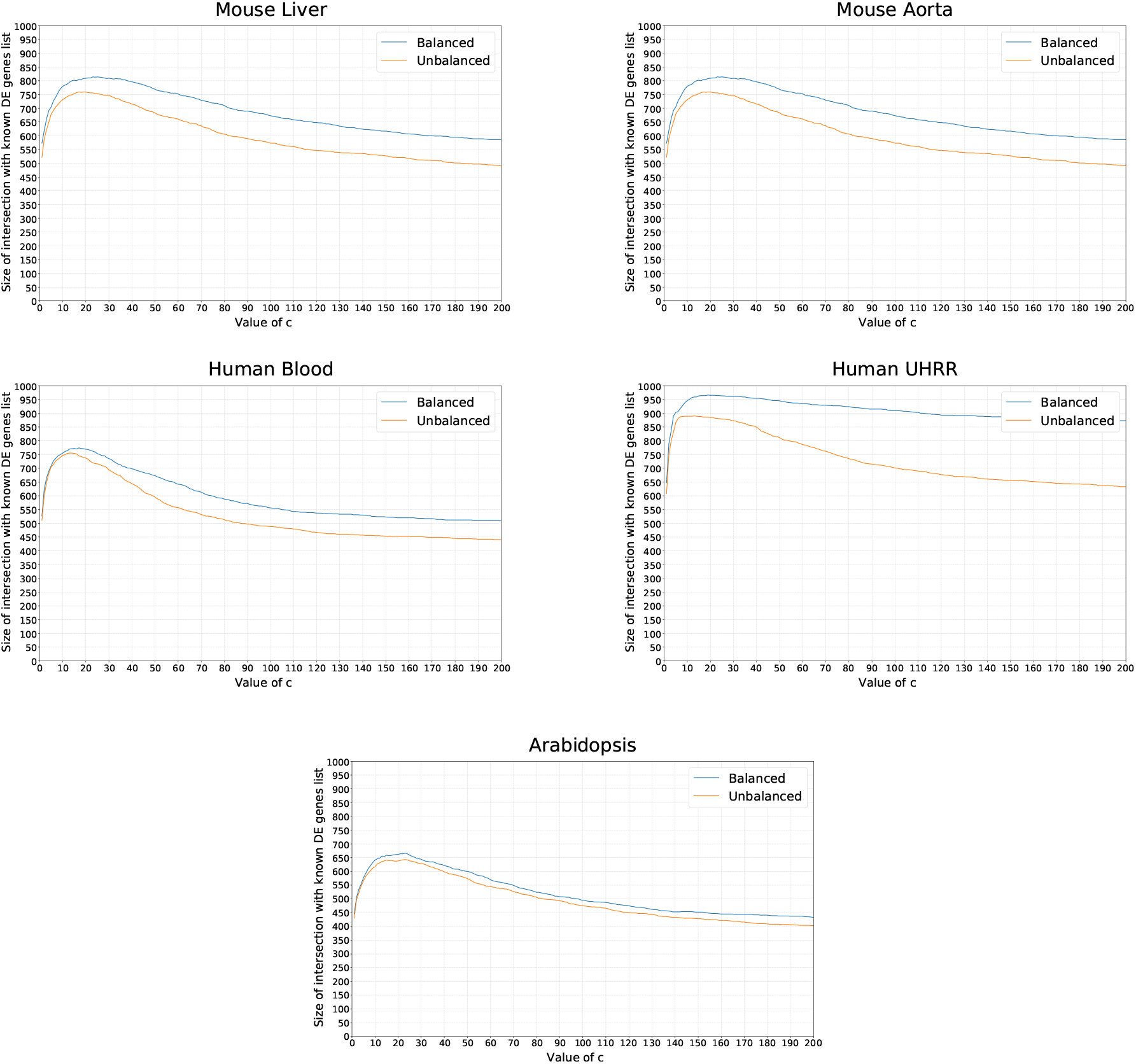
Results for datasets with 1000 DE genes

<u>Observations:</u>

1. In all cases the maximal intersection with the truth occurs around *c* = 20. In fact anything from *c* = 15 to *c* = 25 is close to optimal.
2. The number of DE genes had little effect on the optimal value of c.
3. The widely used value *c* = 1 is extremely sub-optimal.
4. Performance is also extremely sub-optimal for large values of c. Therefore ranking by difference is also a bad idea.

The above results were based on eight replicates in each condition. It is necessary to explore the optimal value of *c* also with fewer replicates (down to as few as one). The results for the 200 DE genes mouse liver dataset (1000 DE genes dataset, respectively) are shown in Fig. 4 (Fig. 5, respectively).

**Figure 4.**
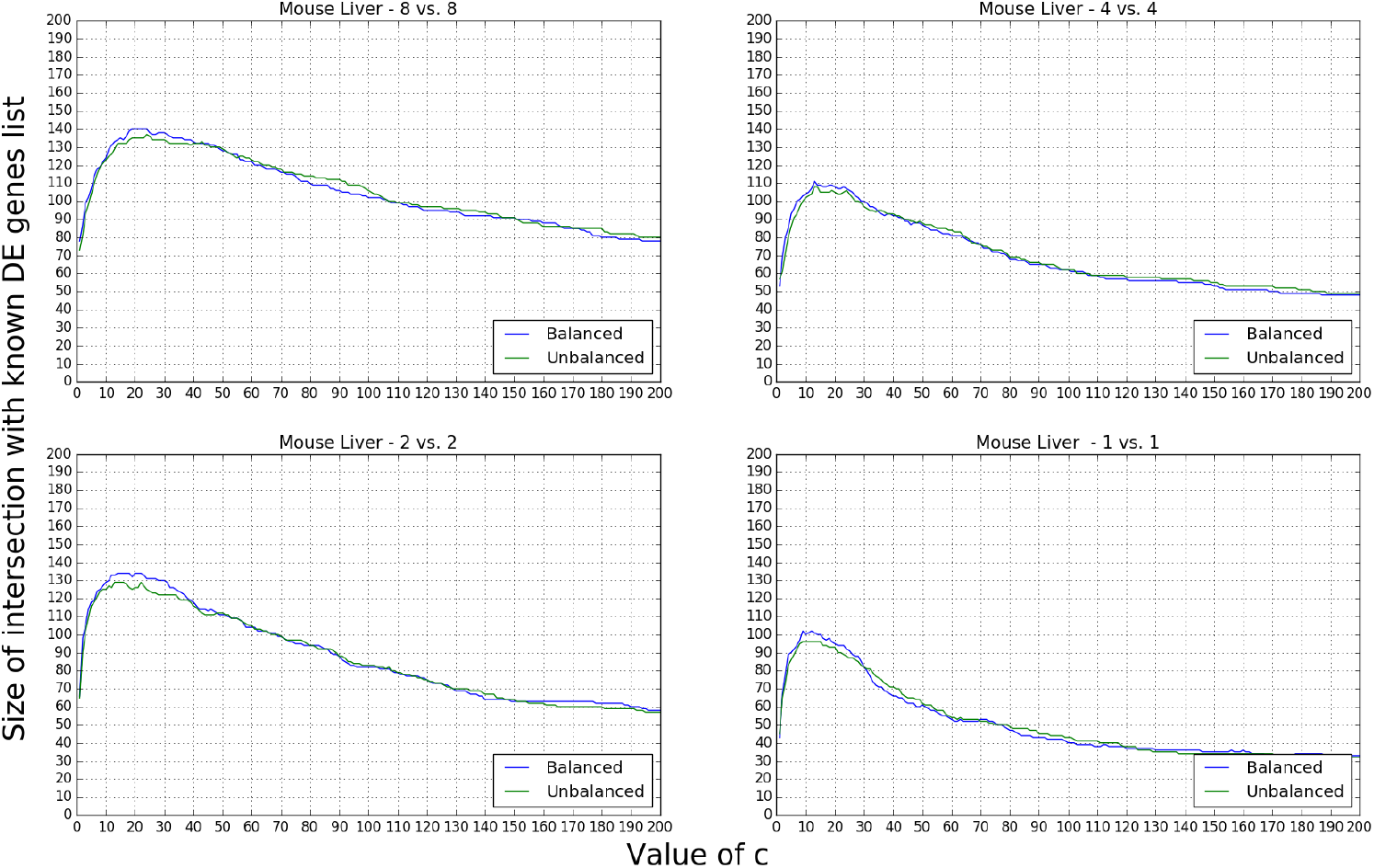
Effect of number of replicates in 200 DE genes dataset (mouse liver)

**Figure 5.**
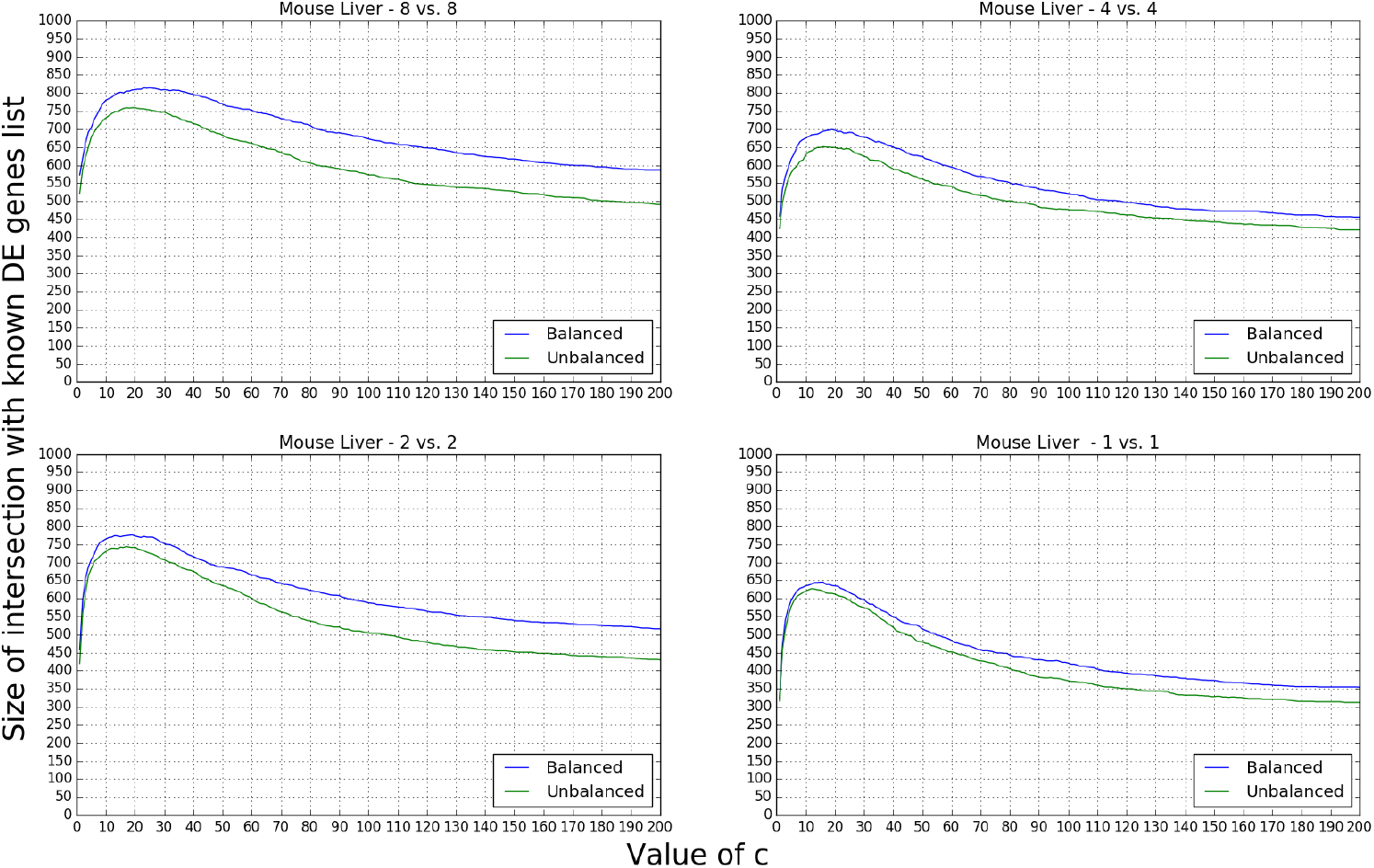
Effect of number of replicates in 1000 DE genes dataset (mouse liver)

**Figure 6.**
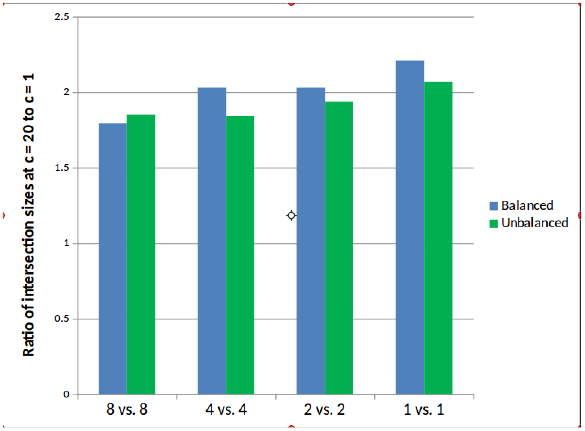
Comparison across number of replicates (200 DE genes dataset)

**Figure 7.**
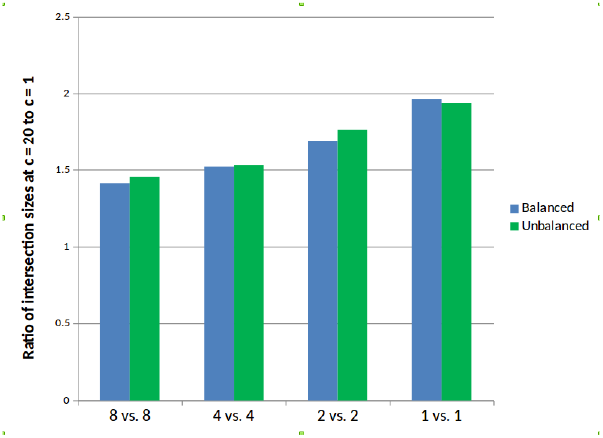
Comparison across number of replicates (1000 DE genes dataset)

<u>Observations:</u>

- Once again, *c* = 1 is much too small. Greatly improved performance is obtained by increasing *c* from 1 to 10. Once again, surprisingly, *c* = 20 is close to optimal in all data sets.
- As can be seen from the figures, the smaller the number of replicates the bigger the improvement by using *c* = 20 instead of *c* = 1.

#### 3.1.1. Application to Pathway Enrichment Analysis

One way to compare two differential expression analyses is by which analysis leads to more meaningful and powerful pathway enrichment *p*-values. In order to evaluate the sensitivity of the pathway enrichment *p*-values on the value of the pseudocount, we employed a RNA-Seq data from a publicly available dataset from GEO repository (accession number - GSE117029). The purpose of the experiment for which the data was generated was to determine whether there is a differential effect on the transcriptome of mice based on time of influenza infection. In order to identify the relevant list of genes in terms of their differential change in gene expression across the time-points, we employed three methods; two-way ANOVA *p*-values, differential change in adjusted fold-change with pseudocount of 1, and with pseudocount of 20. The details of the methods may be found in the supplementary material. The results of the pathway enrichment analysis performed using IPA are shown in Table 1.

**Table 1.** Comparison of pathway enrichment *p*-values based on top 900 genes from the three ranking procedures

| | $-\log p\text{-value}$ | | |
| --- | --- | --- | --- |
|  | 2-Way ANOVA<br>interaction | DC list with<br>pseudocount = 1 | DC list with<br>pseudocount = 20 |
| Ingenuity Canonical Pathways |  |  |  |
| Granulocyte Adhesion and Diapedesis | 4.16 | 14.1 | 17.2 |
| Agranulocyte Adhesion and Diapedesis | 3.32 | 12.4 | 13.6 |
| Role of Pattern Recognition Receptors in Recognition of Bacteria and Viruses | 5.11 | 5.93 | 10.7 |
| Role of Hypercytokinemia/hyperchemokinaemia in the Pathogenesis of Influenza | not significant | 11.2 | 9.66 |
| Communication between Innate and Adaptive Immune Cells | 3.62 | 5.84 | 8.42 |
| Nicotine Degradation II | not significant | 4.48 | 8.1 |
| Differential Regulation of Cytokine Production in Intestinal Epithelial Cells by IL-17A and IL-17F | not significant | 7.58 | 7.84 |
| Activation of IRF by Cytosolic Pattern Recognition Receptors | 7.21 | 3.55 | 4 |

<u>Observations :</u>

1. The enrichment *p*-values are significantly lower in the gene list obtained from adjusted FC with pseudocount as compared to the list obtained by using two-way ANOVA interaction *p*-values.
2. The enrichment *p*-values are generally lower in the case of adjusted FC with pseudocount of 20 compared to adjusted FC with pseudocount of 1. For example, in the case of the pathway “Role of Pattern Recognition Receptors in Recognition of Bacteria and Viruses” the improvement elevates the pathway from being of marginal interest to being quite significant.
3. There are exceptions to the above two observations. For example, the pathway “Activation of IRF by Cytosolic Pattern Recognition Receptors” has significantly better enrichment *p*-values in the list obtained from 2-way ANOVA *p*-values. This could partly be because we use the same number of DE genes (900) for each list and the better enriched pathways tend to have better gene coverage reducing the gene coverage for the less enriched pathways.

### 3.2. Differential Dicodon Usage

The frequency of neighboring amino acids in the proteome varies from species to species. For example valine and arginine are very rarely adjacent in human, however they are very often adjacent in Zika virus. This information can potentially be exploited as a diagnostic because the active translation of any particular di-codon can be detected with FRET signaling using fluorescent tRNAs (Barhoom et al). Towards this goal it is necessary to identify the tRNA-tRNA pairs that have the best chance of providing an informative signature that will reveal the presence of the parasite. In other words, identifying pairs that are high in the virus proteome and low in the human proteome. As using all 1035 possible pairs is not practical with the current state of the technology, we want to find a small handful that work. Therefore we need to rank di-codons by the differential frequencies of the occurrence of tRNA-tRNA anticodon pairs in translating the two proteomes. We want to be able to develop a universal diagnostic that can distinguish between any two viruses and host. They may present similar symptoms (e.g. Zika, Dengue and Chikungunya) and current diagnostics are time-consuming and cumbersome. Also this will allow us to identify whether a virus is latent or active.

As the relative frequencies of most of the tRNA-tRNA pairs are close to 0, the ranking problem based on differential relative frequencies amongst the tRNA-tRNA pairs is a challenging one. This is also a fundamentally different problem from the RNA-Seq example discussed above as there are no replicates and it isn’t amenable to conventional statistical analysis where benchmarking data may be generated with the known ground truth. Since the number of tRNA-tRNA pairs is of the order of magnitude of 1000, *a priori* we expect the relative frequency of each tRNA-tRNA pair to be close to 0.1%. By the very nature of the problem, the appropriate choice of pseudocount in this scenario cannot be directly determined. But from Corollary 2.2 and the discussion following it, a spectrum of values of pseudo-count can be evaluated until satisfactory results are obtained, without having to straitjacket our choice to a value very close to zero. In this case, we appeal to the interpretation mentioned in Main Result 2 that the pseudocount represents the value *a* such that the change 0 → *a* is as significant as a two-fold change for large values. The coding sequences of IVA H1N1 strain (GenBank id - GQ160811.1, Korea 2009) and IVB Victoria strain (GenBank id - CY040455.1, Malaysia 2004) downloaded from the NCBI Nucleotide database were used to obtain the relative frequencies (in percentage) of tRNA-tRNA pairs. Using a pseudocount of 0.5, the adjusted fold-change was computed to rank the tRNA-tRNA pairs for their abilities to distinguish between IVA and IVB infection. In Figure 8(A)-(B), the heatmaps display the relative frequencies of the 1035 tRNA-tRNA pairs in IVA H1N1, IVB Victoria, respectively. The labels are the anticodons associated with the tRNAs. In Figure 8(C)-(D), for the tRNA-tRNA pairs that best distinguish IVA H1N1 and IVB Victoria (called DiDi pairs for the two viruses), the relative frequencies in IVA H1N1, IVB Victoria, respectively, are shown. In Table 2, the relative frequencies of the top DiDi pairs are noted, 3 each for IVA H1N1 and IVB Victoria.

**Figure 8.**
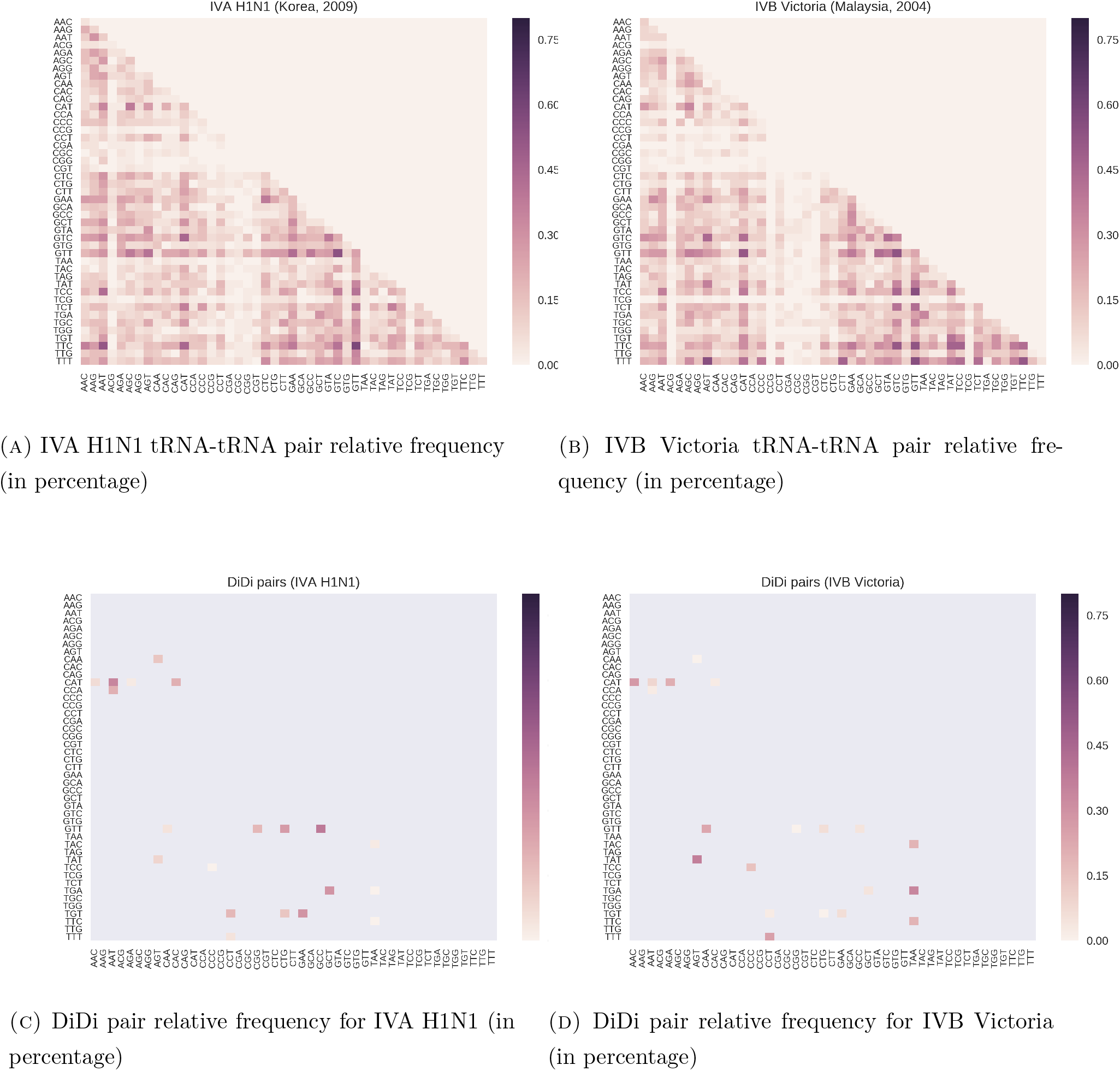

**Table 2.** Top DiDi pairs for IVA H1N1 - IVB Victoria comparison

| tRNA-tRNA pair | relative frequency |  |
| --- | --- | --- |
|  | IVA H1N1 | IVB Victoria |
| TAA-TGA | 0 | 0.340 |
| TAA-TTC | 0 | 0.191 |
| AGA-CAT | 0.022 | 0.212 |
| GCC-GTT | 0.383 | 0.042 |
| CGG-GTT | 0.180 | 0 |
| GCT-TGA | 0.293 | 0.042 |

## 4. Discussion

The procedure of adding a pseudocount of one to all values arises very naturally as a solution to the division-by-zero problem, and certainly thousands of investigators have come to this solution independently. However, it is perhaps counterintuitive to believe that the final results would be highly sensitive to this value. Even so, there have been surprisingly few investigations into this issue. Erhard and Zimmer [6] give an interpretation of pseucocounts as parameters in a prior distibution, when using Bayesian MAP estimation of the fold-change from RNA-Seq, however the focus is on confidence intervals and using pseudocounts tailored to each gene to incorporate prior knowledge. The results were not related to the ranking problem, nor how to optimize the parameters when there is no prior information available. The only other paper we identified which did a systematic benchmarking of pseudocounts is Warden *et al* [2], however they limited the range of their pseudocount to be between 0 and 1; and as we’ve seen the optimal value may be much larger. Furthermore, they did not attempt to develop a general framework in which to investigate these issues. The closest to a generalized framework that has been published is an obscure work by Aczél *et al* [1] which develops a framework in which to compare *p*-values. As *p*-values are between zero and one they focus on the unit square in the 1st quadrant, while an analysis of fold-change involves the entire 1st quadrant.

In [6], the fold-change is discussed as a means of canceling bias which is assumed to grow linearly with the feature counts. The assumption of linearity of bias is deemed more appropriate at the read sequence level during the PCR step in RNA-Seq than the same assumption in terms of read counts at the aggregate features level (such as genes, transcripts, etc.) The authors introduce a count ratio model based on local read counts aligned to a certain genomic position. The parameters used in the prior beta distribution for the log-fold-change ultimately feature in the estimator for the fold-change in the form of pseudocounts. The total fold-change of a feature is modeled as a weighted average of local fold-changes, where positions with many reads contribute more to the total fold-change than positions with fewer reads.

A similar problem arises in ranking features by *T*-statistic. Let x = (*x*_1_,…, *x_n_*), y = (*y*_1_,…, *y_n_*) be two vectors in 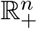. The *T*-statistic is given by a function 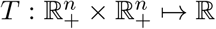 which sends (**x, y**) to 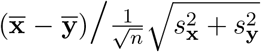 where the denominator denotes the pooled standard deviation. In this case, problems arise when the variance of both groups is small. Firstly since the denominator is essentially given by these variances, that can cause the *T*-statistic to blow up just like dividing by very small numbers in the fold-change. This naturally leads to a higher dimensional analog of the problem. Instead of iso-relevance functions of one variable, addressing these issues will require generalizing this concept to *n* dimensions. Secondly, if a hypothetical dataset consists of observations with zero variance (or all with the same variance), it essentially reduces to the onedimensional problem of assessing relevance of change of real values. The standard procedure of sorting features according to *p*-values (from the *T*-statistic) reduces to sorting by difference which, as discussed earlier, is more convincing at smaller scales but not so much at larger scales. Thus one should keep in mind that any generalization to higher dimensions should be compatible with the concept of iso-relevance functions in a lower dimensional space. Thirdly, the use of the *T*-test implicitly assumes that the variance of the expression of each gene is the same. When the scale of the expression level is different, the variance may be significantly different. To address this issue, similar to using pseudocounts, Huber *et al* introduced what they called a “fudge factor” added to the denominator. The rankings of features can be remarkably sensitive to this parameter (see Grant *et al* [4]). This will be the focus of future work.

We point out another fundamental difference between the usage of ranking by fold-changes vs. *p*-values with regards to pathway enrichment analyses. The *p*-values measure our confidence in the fact that the corresponding genes are differentially expressed. They convey little information about the ‘effect size’ or relevance of the change. Although a *p*-value, or more precisely, FDR (false discovery rate) cutoff works well for determining the pool of DE genes but it is by nature quite ineffective in actually sorting the DE genes by effect size. There are two issues here which deserve further investigation. First, for the purpose of pathway enrichment analysis although the actual ranking may not be as important, sometimes we may care about the relative rankings of certain pathways. Second, often finding the right FDR cut-off for pathway enrichment analysis is an issue of trial-and-error and preliminary investigations have shown that this phenomenon may be because we do not take the effect size into account for sorting. We will leave that discussion for another occasion.

## 5. Materials and Methods

### Benchmarking Data

The RNA-Seq data was obtained from GEO datasets (GSE95802, GSE115264, GSE99719) and manipulated into benchmarking data. For each data set we started with 16 samples from two conditions. Samples were permuted to have two new groups with 8 samples from each condition in each group. The distribution of *p*-values was then plotted to make sure it does not deviate significanty from uniform. From this data, 200 genes were chosen from each of 10 (average) expression levels low to high. For each gene chosen, reads were randomly removed from the samples in one group to mimic differential expression. 10 different levels of differential expression were created from 1.1 fold to infinite fold. This was done in two ways, one way all 200 genes are up-regulated; in the second way half are upregulated and half are downregulated. Another set of data sets was generated similarly, however with 1000 DE genes instead of 200. Benchmarking data is described in more detail and available for download at http://bioinf.itmat.upenn.edu/benchmarking/rnaseq/port/datasets.php

### Mouse Influenza data

The RNA-Seq data was obtained from a GEO dataset (GSE117029) for an experiment involving male mice infected at two distinct time points (three replicates at 6 a.m., four replicates at 6 p.m.) with influenza virus and three replicates each from PBS mock infected controls at the same two time points. The purpose of the experiment was to determine whether there is a differential effect on the transcriptome of mice based on time of infection. As several genes are differentially expressed simply by virtue of the circadian rhythm or other time-dependent effects, it was necessary to account for this in the analysis. A standard approach is to perform 2-way ANOVA with two categorical variables: “treatment status” (control or treated) and “time” (6am or 6pm) and to look for genes with significant interaction *p*-values. Although it is generally considered suboptimal with just three replicates per condition, for the sake of comparison we performed a two-way ANOVA analysis to find interaction effects and sorted the list of genes by the interaction *p*-values. At each timepoint the adjusted fold change of gene expression level was computed relative to the corresponding control group using a pseudocount of 20. Genes which had coefficient of variation > 3 were filtered out and the remaining genes were then ranked based on the ratios of their adjusted fold-change at the two timepoints. Out of a total of 20401 genes, we found 900 genes exhibiting a differential log fold change (DC) of atleast 0.67 in magnitude which corresponds to about 5-fold change in the adjusted fold-changes. We repeated the above ranking procedure for the list of genes using a pseudocount of 1. For a consistent comparison, we noted the top 900 genes in all three of the ranking procedures to check for pathway enrichment.

### Di-Codon data

The coding sequences for IVA H1N1 strain (Korea - 2009), IVB Victoria strain (Malaysia - 2004) were downloaded from the NCBI Nucleotide database using the GenBank IDs GQ160811.1, CY040455.1, respectively. The di-codon frequency in the CDS’s of both viruses was computed. The final goal of this exercise, which is out of scope of the paper, was to identify DiDi pairs which have low usage frequency in the human ORFeome as that would be the background in any diagnostic procedure. As viruses use the host’s translational machinery, in order use DiCoMPS ([5]) one needs to reinterpret dicodon usage in terms of tRNA-tRNA pair usage. In general there is a fairly strong correlation between relative frequency of codon usage and the number of tRNA genes having an anticodon sequence complementary to the codon. For the common 20 amino acids, there are 55 isoacceptor tRNAs that are complementary to the 61 codons. The 6 isoacceptors (GCC, AGT, ACG, AAG, GAA, GTG) that completely decode 2 codons each, have an *A* or *G* at the 5’-end of the anticodon sequence that forms a base pair with either a *U* or *C* at the 3’-end of the codon - the wobble position. In addition there are 8 pairs of isoacceptor tRNAs (see supplementary Table 1, colored in yellow), that differ by having an *A* or *G* at the 5’-end of the anticodon sequence that have very different numbers of tRNA genes (8 - 38fold differences) but are complementary to codons with similar relative frequency of codon usages (http://gtrnadb.ucsc.edu/GtRNAdb2/genomes/eukaryota/Hsapi38/Hsapi38-summary-codon.html). The clear inference is that in these cases the dominant tRNA isoacceptor will generally translate both codons. Finally, there are 2 pairs of anticodons (see supplementary Table 1, colored in blue), again differing by having an *A* or *G* at the 5-end of the anticodon sequence, where the difference in the number of tRNA genes is less dramatic (3 6-fold differences) and are complementary to codons with similar frequency of codon usages. In these cases it is likely that both tRNAs will be involved in translating the two codons, but again the dominant tRNA isoacceptor is likely to play the major role. Based on the above reasoning, we focussed on the 45(= 61 – (6 + 8 + 2)) tRNA isoacceptors that are most strongly involved in translation. With this mapping of codons to tRNA isoacceptors, the relative frequencies of each of the 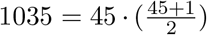 tRNA-tRNA anticodon pairs were computed.

## References

[1] J. Aczél, G. Rote, J. Schwaiger; Webs, Iteration Groups and Equivalent Changes in Probabilities, Quarterly of Applied Mathematics 54 (1996), 475–499.

[2] Warden CD, Yuan YC, Wu X.; (2013) Optimal Calculation of RNA-Seq Fold-Change Values. International Journal of Computational Bioinformatics and In Silico Modeling 2(6), 285–292.

[3] Tusher V, Tibshirani R, Chu C (2001). Significance analysis of microarrays applied to transcriptional responses to ionizing radiation, Proceedings of the National Academy of Sciences 98:51165121.

[4] Grant GR, Liu J, Stoeckert C (2005). A practical false discovery rate approach to identifying patterns of differential expression in microarray data.

[5] Barhoom S, Farrell I, Shai B, Dahary D, Cooperman BS, Smilansky Z, Elroy-Stein O, Ehrlich M. Dicodon monitoring of protein synthesis (DiCoMPS) reveals levels of synthesis of a viral protein in single cells. Nucleic Acids Res. 2013 Oct;41(18):e177.

[6] Erhard F, Zimmer R. Count ratio model reveals bias affecting NGS fold changes. Nucleic Acids Res. 2015 Vol. 43. No.20.

[7] Michael Love, Wolfgang Huber, Simon Anders. Moderated estimation of fold change and dispersion for RNA-seq data with DESeq2. Love et al. Genome Biology (2014) 15:550 DOI 10.1186/s13059-014-0550-8

